# An Array of 60,000 Antibodies for Proteome-Scale Antibody Generation and Target Discovery

**DOI:** 10.1101/553339

## Abstract

Antibodies are essential for elucidating the roles of genes decoded by genome sequencing. However, affordable technology for proteome-scale antibody generation does not exist. To address this, we developed the Proteome Epitope Tag Antibody Library (PETAL) and its array. PETAL consists of 62,208 mAbs against 15,199 peptides from diverse proteomes. PETAL harbors binders for a great multitude of proteins in nature due to antibody multispecificity, an intrinsic feature of an antibody. Distinctive combinations of 10,000-20,000 mAbs were found to target specific proteomes by array screening. Phenotype-specific mAb-target pairs were discovered for maize and zebrafish samples. Immunofluorescence and flow cytometry mAbs for human membrane proteins and ChIP-seq mAbs for transcription factors were identified from respective proteome-binding PETAL mAbs. Differential screening of cell surface proteomes of tumor and normal tissues discovered internalizing tumor antigens for antibody-drug conjugates. By discovering high affinity mAbs at a fraction of current time and cost, PETAL enables proteome-scale antibody generation and target discovery.

## INTRODUCTION

Facilitated by the ever-growing capability of current DNA sequencing technologies, more than 1,300 genomes of animals and 496 genomes of plants and many others have already been sequenced, representing millions of genes, and the number will increase faster from projects such as G10K, i5k and so on^1^. To understand the roles of these genes, the functions of the gene-coded proteins need to be explored, and antibodies, especially renewable monoclonal antibodies (mAbs) generated at a proteome-scale, are urgently needed. mAbs produced by hybridoma technologies for human proteins have long been recognized as the most direct tools for diagnostic and therapeutic target discovery^2^. Classic therapeutic targets Sialyl Lewis Y, prostate-specific membrane antigen (PSMA) and, more recently, a novel target for multiple myeloma were discovered by mAbs for cell surface proteins^3–6^.

Despite the power of mAbs and mAb-based discovery, large-scale generation of mAbs remains difficult since traditional hybridoma development is time-consuming (4-6 months starting from antigens), expensive ($3,000-8,000 per antigen), and challenging to scale. Further, mAb generation by immunization typically requires a milligram of purified antigens, a significant challenge for many protein targets, especially for membrane proteins of primary research interest. Even for human proteins, a majority of the 6,000 membrane proteins have not been directly explored as diagnostic or therapeutic targets due to a lack of high-quality antibodies for applications such as flow cytometry (FACS) and immunofluorescence (IF)^7, 8^

The Human Protein Atlas (HPA) offers an alternative approach for proteome-scale antibody development. HPA has generated over 25,000 affinity-purified polyclonal antibodies against >17,000 human proteins covering more than 80% of the human proteome^9, 10^ However, it is impractical to replicate the success of HPA on a significant number of other species with a need for proteome-scale antibodies due to the great human and capital resources required for such a project. Furthermore, polyclonal HPA antibodies are not renewable, making reproduction of these antibodies with consistent quality difficult. Thus, proteome-scale antibody generation has remained elusive for most sequenced genomes.

Over the past few decades, numerous attempts have been made to address high cost and poor scalability of large-scale antibody generation by improving traditional hybridoma methods and developing better *in vitro* recombinant antibody libraries and more efficient screening technologies^11–14^. For *in vitro* methods, continuing development of novel display technologies^15–17^ and improvements in library design and screening methods^18^ have been attempted dating back twenty years. However, established synthetic antibody libraries for therapeutic antibody discovery aren’t yet used for large-scale reagent generation due to the concern of high cost. Despite an attempt to generate research antibodies using phage display libraries^19^, it is not economical to use these resources for generating antibodies for nonclinical uses or for nonhuman proteomes.

Antibody microarray is a powerful platform for high-throughput, multiplexed protein profiling using a collection of immobilized antibodies^20, 21^. By using antibody array, one can achieve low cost and fast antibody discovery by direct array screening. In one approach, a library of ~10,000 in silico-designed and – synthesized antibody fragments was used to build an antibody array for *de novo* antibody discovery^22^. The arrayed library was able to generate antibody leads with μM binding affinity for therapeutic protein targets, suggesting a spatially addressed library comprising tens of thousands of individual antibodies should be sufficient for antibody discovery. However, this synthetic antibody library screening approach did not achieve broader impact since required additional antibody affinity maturation and engineering limited its usefulness for routine research affinity reagent development and target profiling.

Here, we present a system integrating “¡ndustrial-scale” hybridoma development, antibody microarray, and affinity proteomics to overcome previous challenges for proteome-scale antibody development and target discovery. Our technology, called the Proteome Epitope Tag Antibody Library (PETAL), takes advantage of antibody multispecificity^23–25^, an intrinsic property of antibody molecules to bind to a large number of proteins unrelated to the original antigen that the antibody was raised against with high affinity and specificity. Antibody multispecificity is exemplified by an anti-p24 (HIV-1) peptide monoclonal antibody. Its epitope sequences, consisting of key interacting residues deduced from five unrelated peptides binding to CB4-1, were identified in hundreds of heterologous proteins, and those proteins that could be obtained were shown to bind CB4-1^26^. PETAL is a mouse monoclonal antibody library consisting of 62,208 mAbs made against 15,199 peptide antigens representative of 3,694 proteins from 418 proteomes. PETAL has the potential to bind to a large number of proteins in nature due to antibody multispecificity. An antibody microarray was fabricated using PETAL mAbs. Using cell lysates of diverse proteomes to screen the PETAL array, we demonstrated, for the first time, the feasibility for proteome-scale protein targeting by mAbs. Identified antibodies are capable of broad applications, as shown for human membrane and nuclear protein targets. Phenotype-specific mAb-target pairs were identified for maize and zebrafish from respective proteome-specific mAbs. Therapeutic target candidate discovery was demonstrated by differential screening of normal versus tumor membrane proteomes. An antibody targeting ProteinX was identified as a candidate for building an antibody-drug conjugate (ADC) for Lung Squamous Cell Carcinoma (LUSCC) *in vitro* and *in vivo.* By generating high affinity mAbs at a small fraction of current time and cost, PETAL enables affinity reagent generation and target discovery for proteomes with available genomic sequence information.

## RESULTS

The immunological basis of PETAL is antibody multispecificity (**Fig. 1a**). PETAL is an antipeptide antibody library of 62,208 monoclonal antibodies designed to have the potential for harboring binders for a great number of proteins from diverse proteomes (**Fig. 1b**). When PETAL is immobilized in an array format (**Fig. 1c**), it enables proteome-scale antibody generation and differential target discovery (**Fig. 1d**).

**Figure 1.**
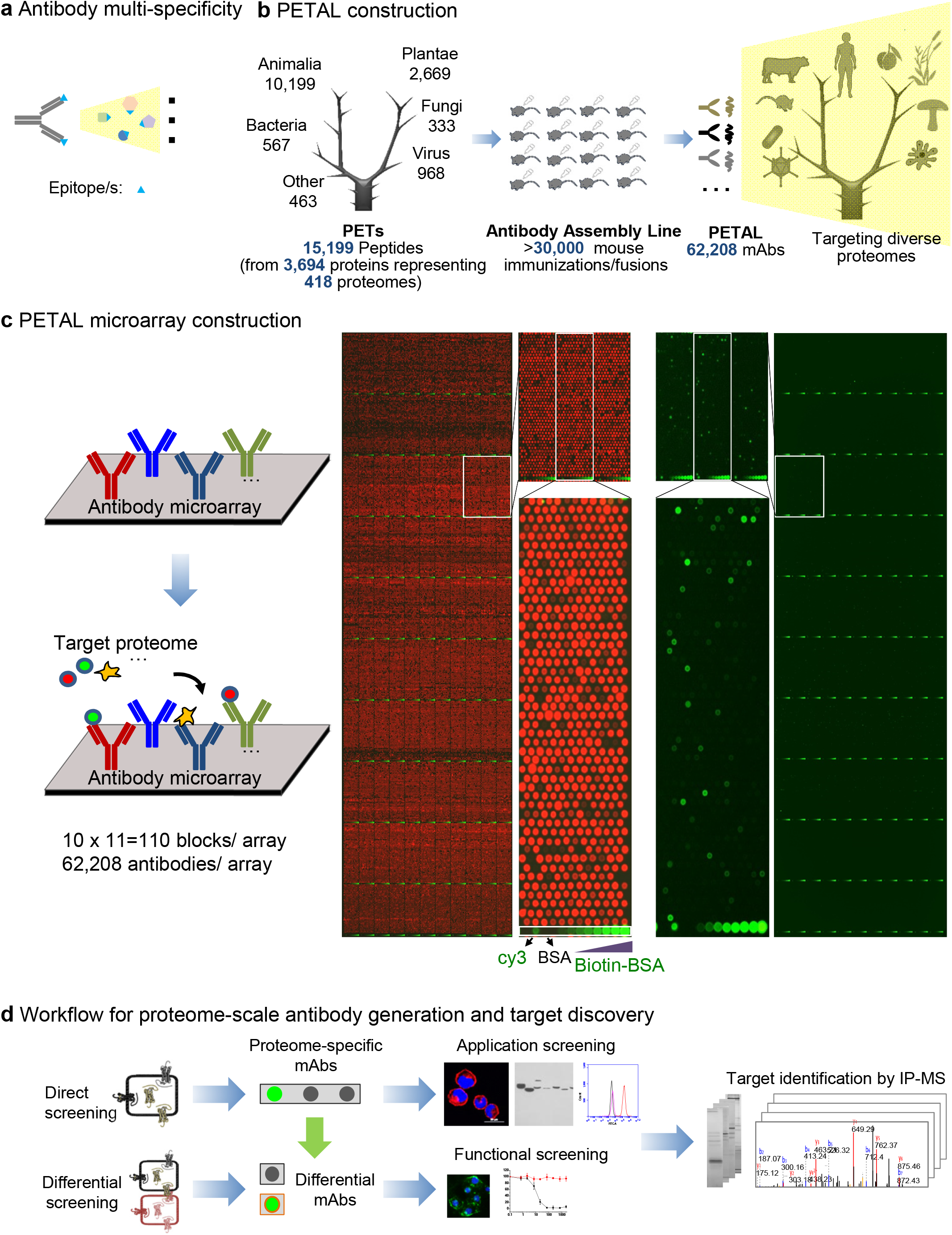
Construction and application of PETAL and its array for antibody/target discovery. (a) Antibody multispecificity. An antibody binds to an epitope/mimotope found within a variety of proteins from different species, leading to high affinity, specific binding of this antibody to a large number of proteins in nature. (b) PETAL construction. PETAL is a library of 62,208 mouse monoclonal antibodies (mAbs) derived from immunization of more than 30,000 mice against 15,199 diverse peptide antigens. PETAL has the potential to recognize a great number of proteins in nature. (c) PETAL microarray construction. PETAL is printed into an antibody microarray as a high-throughput platform for antibody/target discovery (left panel). Right panel shows the design/layout of the array (red, visualized by a Cy5-conjugated anti-mouse antibody) and an array hybridization result using a protein sample (positive-binding mAb spots are shown as green). (d) Workflow for proteome-scale antibody generation and target discovery. Two array screening applications are shown: direct screening to identify proteome-specific mAbs and subsequent antibody application screening and target identification or differential screening to discover mAbs and their cellular targets associated with a specific phenotype. Differential screening can also be performed with proteomic-specific mAbs (light green arrow) as in the examples of maize and zebrafish data.

### Design and construction of PETAL

A total of 15,199 peptide antigens, called Proteome Epitope Tags or PETs, were designed from 3,694 proteins representing 418 proteomes (**Fig. 1b; Supplementary Fig. 1a**). Within each proteome, PETs were selected from unique regions of protein sequence using the heuristic blastp algorithms optimal for short peptide sequence comparison^27, 28^ Sequence analysis showed PET sequences were random and diverse (**Supplementary Fig. 1b**).

To construct PETAL, PET antigens were synthesized and used to generate mouse mAbs by a large-scale monoclonal antibody development operation modeled after an assembly line (**Fig. 1b**). More than 30,000 mice were immunized at an average of 2 mice per PET. A total of 62,208 mouse monoclonal antibodies were generated. Each hybridoma cell line was used to prepare ascites containing 1-20 mg of mouse IgGs with varying concentrations from 0.1 to 10 mg/mL; most were in the 1-3 mg/mL range.

### PETAL diversity

To evaluate the diversity of PETAL, particularly multiple mAbs generated using the same PET, antibody V-region was sequenced for 91 randomly selected hybridomas, including 68 hybridoma clones against 24 PETs with 2-5 mAbs/PET and 23 hybridomas against 23 unique PETs (**Supplementary Fig. 1c-e**). Sequences with ≥2 amino acid differences in the CDR regions were considered “unique”, although two antibodies with a single CDR amino acid difference were found to bind to different epitopes^29^. Close to 90% of sequences were unique (**Supplementary Fig. 1d**). Multiple mAbs generated against the same PET peptide antigen were mostly (80-100%) unique (**Supplementary Fig. 1e**), indicating the effective library size was close the total number of PETAL mAbs since framework differences could also contribute to different binding affinity and therefore specificity.

### Construction of PETAL microarray with 62,208 mAbs

To use PETAL for multiplexed screening, an antibody microarray was constructed with all 62,208 antibodies (**Fig. 1c**). Array printing quality was assessed using an anti-mouse secondary antibody conjugated with Cy5 (**Fig. 1c**, left two panels). Antibodies of 10-1000 pg per spot (mostly 100-300 pg) were detected with a fluorescent intensity ranging from 500 to >60,000 (**Supplementary Fig. 1f**).

Typically, biotinylated antigens such as 1-10 mg of proteome samples (**Fig. 1c**, right two panels) were used for array screening. Array screening consistency was investigated using a human plasma sample in three independent replicates. An average of 20,000 antibodies showed positive binding ((signal intensity-background)/background > 3) with a coefficient of variation (CV) of 7%. More than 90% of array-positive antibodies had a fluorescent intensity of 500-10,000. The Pearson’s correlation coefficient (r) among triplicate experiments was 0.98 (**Supplementary Fig. 1g**). These data establish the PETAL array as a reproducible platform for screening.

To test the library/array for generating mAbs, a total of 81 recombinant proteins were used to screen the PETAL array (**Supplementary Fig. 2, Table S3**). Approximately half (47%, 38/81) of the proteins produced an average of 3.7 (141/38) mAbs/antigen after ELISA screening of array binding mAbs using a detection limit of ≤1μg/mL of antigen. When tested in immunoblotting assays, 31% (25/81) and 26% (21/81) of targets were successful on recombinant proteins and endogenous samples, respectively (**Supplementary Fig. 2, Table S3**).

### PETAL targets diverse proteomes for antibody and target discovery

To apply PETAL for targeting broad and diverse proteomes, 11 proteome samples of plants, animals and bacteria were used to probe PETAL arrays (**Fig. 2**). The number of array-positive antibodies for each proteome was 10,000-20,000. A selection of ~1,000/proteome of array-positive antibodies across the fluorescent intensity scale was used to probe endogenous samples to estimate the number of antibodies suitable for immunoblotting (**Fig. 2a**). Typically, 20-30% of antibodies produced specific (single or predominant single band) immunoblotting results, giving a proteome of 2,000-6,000 potential immunoblotting mAbs. Although proteome-binding mAbs overlapped significantly among proteomes (30-60%), the same antibodies often specifically recognized targets of different size in different proteomes (**Fig. 2b**), likely to be unrelated proteins.

**Figure 2.**
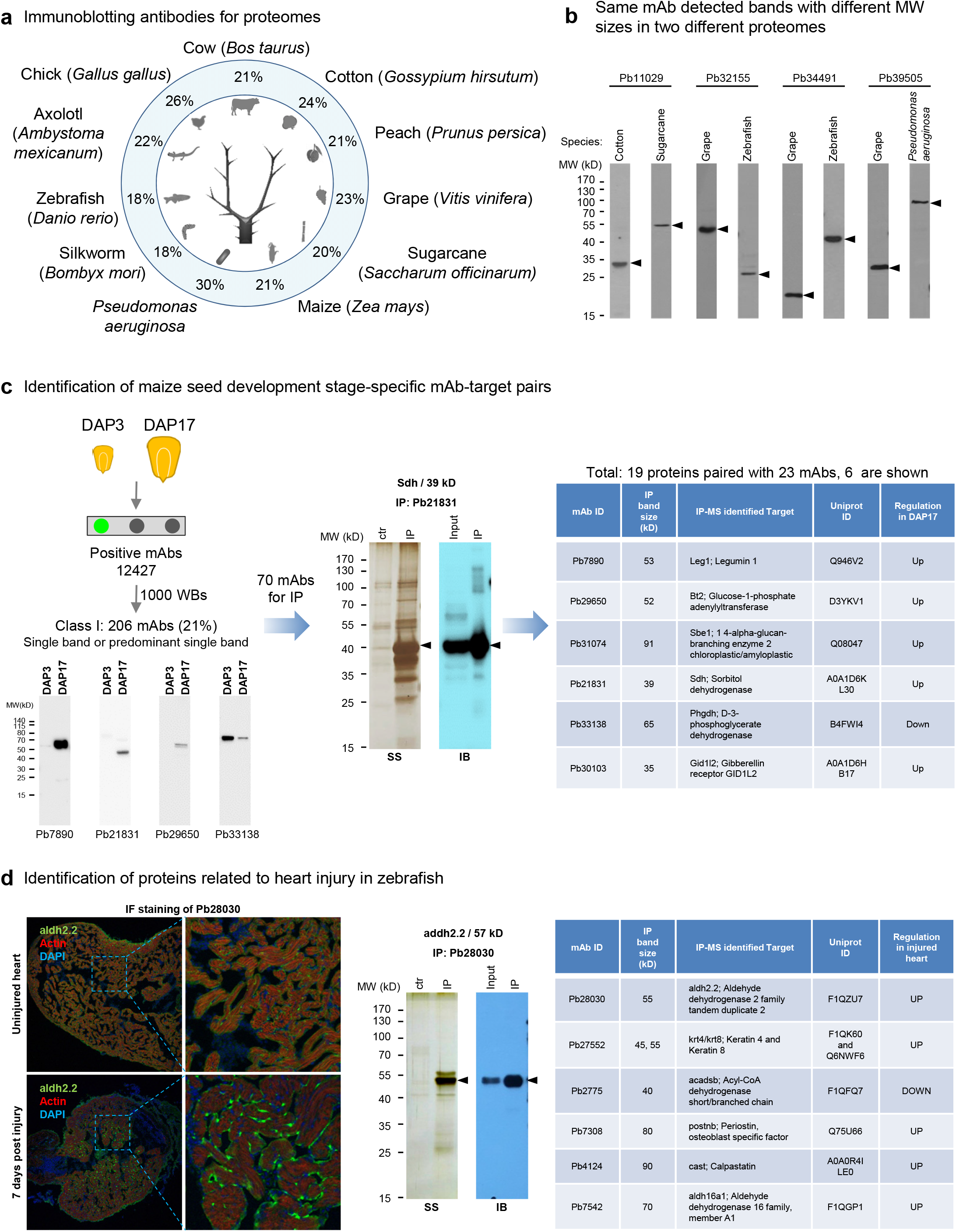
PETAL targets diverse proteomes for antibody and target discovery. (a) Proteome-targeting PETAL mAbs for immunoblotting. Successful rates (labeled as %) of immunoblotting (producing single/predominant single bands) were shown for 11 organisms by using a panel of ~1,000 proteome-binding mAbs for each organism to probe proteome samples. The total number of binding mAbs for a proteome was in the range of 10,000-20,000. The specific tissues for immunoblotting were cow (breast, ovary, and liver), cotton (ovule), peach (leaf or fruit), grape (nuclear fraction of seed), sugarcane (stalk), maize (seed), *Pseudomonas aeruginosa* (whole cell), silkworm (larva), zebrafish (embryo, heart or other tissues), axolotl (regenerating limb), chick (cell lines and tissues). (b) Different targets detected by the same mAb in two different proteomes. Four example antibodies each recognized a specific band with different size in different proteomes upon immunoblotting. (c) Identification of maize seed development stage-specific mAb-target pairs. PETAL array screening using a proteome sample consisting of total protein extracts from maize seeds DAP3 and DAP17 produced a total of 12,427 binding mAbs. A selection of 1,000 mAbs to probe DAP3 and DAP17 samples yielded 206 Class I mAbs (single or predominant single band) and additional 129 Class II mAbs (multiple bands) upon immunoblotting (**Supporting data for Fig. 2**). Seventy differentially expressed mAbs between DAP3 and DAP17 were used to IP their cellular protein targets for MS analysis, resulting in the identification of 19 proteins paired with 23 mAbs. Six proteins are shown in the right panel. Gel pictures from left to right: Class I mAb immunoblotting examples: silver staining (SS) of IP products and immunoblotting (IB) of input and IP products for Pb21831. (d) Identification of heart injury-related proteins from zebrafish. From left to right: IF staining, silver staining of IP products and immunoblotting of input and IP products for Pb28030, and summary of the identified protein targets.

Proteome-specific mAbs were used to identify phenotype-specific mAbs and corresponding protein targets for maize seed development and zebrafish regeneration. A panel of 335 immunoblotting-positive mAbs were produced from 1000 array-positive binding mAbs using a protein sample consisting of maize seeds at two developmental stages, day after pollination 3 (DAP3) and DAP17 (**Fig. 2c**)^30^. These included 206 mAbs with single or predominant single bands (Class I, 21%, 206/1000) and an additional 129 mAbs with more than one (typically 2-3) dominant bands (Class II) on immunoblotting (**Fig. 2c**, **Supporting data for Fig. 2**). After probing DAP3 and DAP17 protein samples separately using Class I mAbs, a panel of 70 differentially expressed mAbs were selected for target identification by immunoprecipitation (IP) and liquid chromatography-tandem mass spectrometry (LC-MS/MS). Nineteen protein targets paired with 23 mAbs were identified, and six examples are shown in **Fig. 2c**, including Sdh, Bt2 and Sbe1, which have previously been shown to be involved in maize seed development^30, 31^.

In another example, a panel of 45 binding mAbs identified from the screening of protein lysates of zebrafish heart was further characterized by IF assays using samples of heart before injury and 7 days post injury (**Fig. 2d**). Six of them showed up-or downregulation in injured heart. Protein targets of these six antibodies were identified by IP and LC-MS/MS (**Fig. 2d**, right panel), including proteins with known heart injury protection function (for example, aldh2.2 in **Fig. 2d**, left and middle panels)^32^.

### Proteome-scale antibody generation for human membrane and nuclear proteins

To apply PETAL for antibody generation for organelle proteomes, PETAL arrays were screened with total protein extracts of membrane and nucleus from human PC9, HepG2, THP-1 and Jurkat cell lines (**Fig. 3a**). The total number of positive mAbs for each sample was in the range of 12,000-18,000.

**Figure 3.**
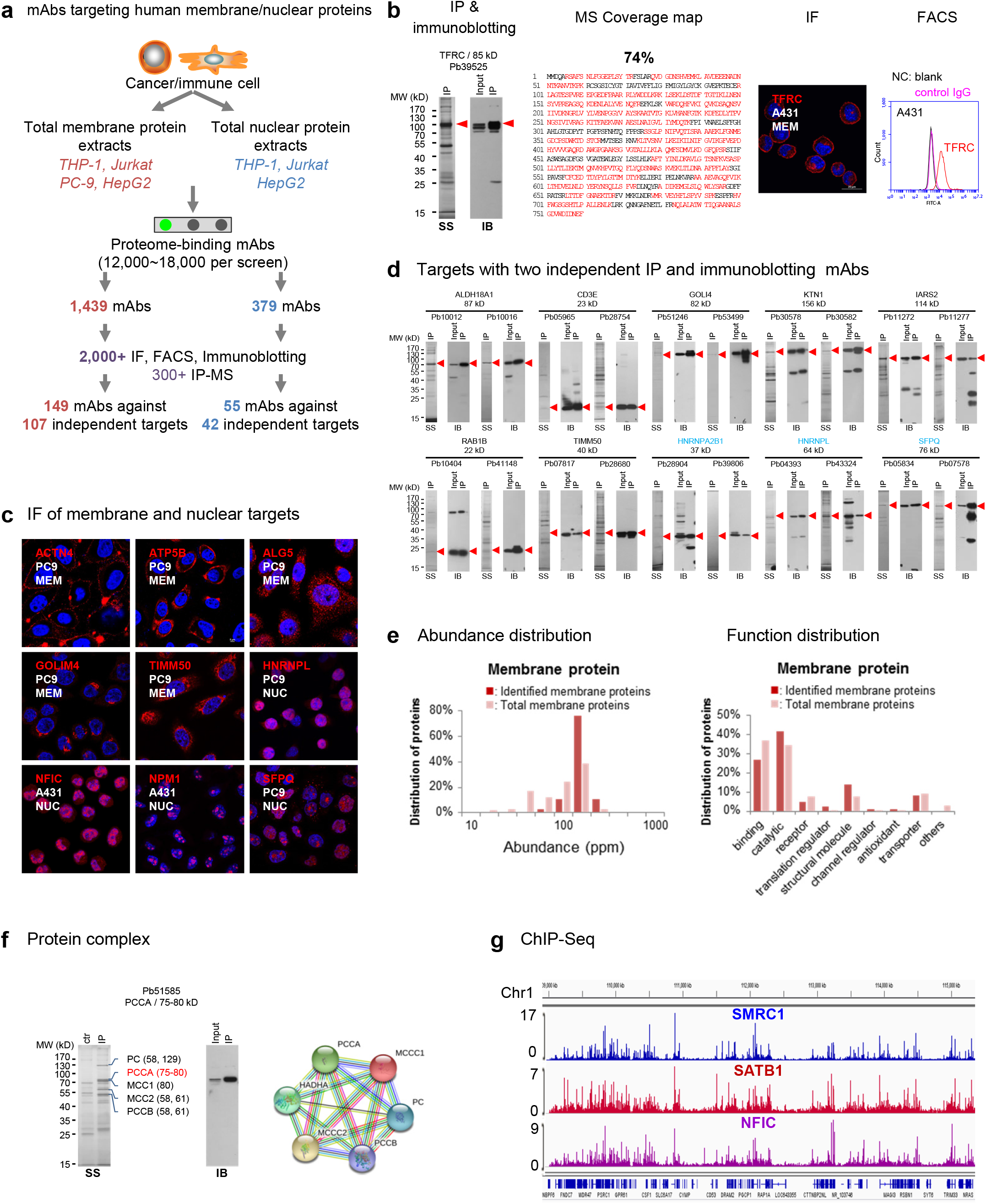
Proteome-scale antibody generation for human membrane and nuclear proteomes. (a) Protein target identification for human membrane and nuclear proteome-specific PETAL mAbs. (b) An example (TFRC/CD71) for antibody target identification. From left to right: silver staining (SS) for IP product, immunoblotting blot (IB) of input and after IP samples, coverage map of MS-identified peptides, IF and FACS data. The cell line for IF was selected according to HPA data. Membrane or nuclear targets were labeled MEM or NUC. Negative controls for FACS included staining with blank and irrelevant IgG. (c) Examples of IF data for endogenously expressed membrane and nuclear proteins. ACTN4 and ACTP5B were stained under nonpermeable conditions. Other targets were stained under permeable staining conditions. (Also see **Movies S1-S6**.) (d) Targets with two independent IP and immunoblotting mAbs. Panel label (SS and IB) was the same as in (b). Nuclear targets were labeled in blue. (e) Abundance and function distribution of proteins identified from the Jurkat cell membrane proteome. (f) Protein interactome example using Pb51585 against PCCA. Protein-protein interacting map (right) analyzed by STRING with the mass-identified proteins. (g) Snapshot of the Integrative Genomics Viewer (IGV) showing sequencing read density of ChIP-seq data generated with antibodies against SMRC1, SATB1 and NFIC in HepG2 cells.

To further screen for application-specific antibodies and to identify their cellular targets, array positive antibodies with high (>10,000), medium (2,000-10,000) and low (500-2,000) fluorescent intensity (**Supplementary Fig. 3a**) were selected. A total of 1,439 positive antibodies for membrane proteomes and 379 for nuclear proteomes were subjected to immunoblotting, IF/FACS and IP assays (**Fig. 3b and c, Supplementary Fig. 3b, Movies S1-S6, Supporting data for Fig. 3**) and their cellular targets were identified by LC-MS/MS (**Fig. 3b**). A total of 149 antibodies representing 107 proteins were identified from membrane proteome screening (**Tables S4 and S5**), including known CD and RAB (small GTPase) molecules CD3e, CD49d, CD71 (TFRC), CD222, CD5, CD2, PROTEINX, RAB1B and RAB14. For nuclear proteomes, a total of 55 antibodies representing 42 proteins were identified (**Tables S4 and S5**), including transcriptional regulators NONO, NFIC, TRIM28, CSNK2A1, MTA2, SATB1, SFPQ, and SMARCC1. Approximately 20% of the targets had at least two independent antibodies that yielded similar IP and immunoblotting results, strengthening the antibody validation quality^33, 34^ (**Fig. 3d**). The success rate of target identification was consistent over a wide fluorescent intensity range (**Supplementary Fig. 3a**), suggesting that more than 1,000 proteins could be covered with 12,000-18,000 proteome-binding antibodies.

Only 20-30% of the array-positive antibodies were successful in immunoblotting assays likely due to the native protein states in the screening samples. For antibodies without successful immunoblotting data (size unknown), overexpression or knock-down experiments were necessary to determine their cellular targets after immunoprecipitation (IP) and mass-spectrum (MS). For example, recognition of an antibody (Pb2795) to a multi-pass ion channel, PIEZO1^35^, was confirmed by the colocalization of IF staining signals with overexpressed PIEZO1-GFP (**Supplementary Fig. 3c**).

To investigate if proteins targeted by antibodies were biased toward specific types, protein abundance and function class (GO annotations) were examined for 87 Jurkat proteins (67 membrane and 20 nuclear) (**Fig. 3e, Supplementary Fig. 3d**). Abundance distribution of these proteins was similar to that of total membrane or nuclear proteins in Jurkat cells according to the PaxDb database^36^. Proteins identified by antibodies were from diverse protein families and functional groups similar to total human membrane and nuclear proteins (**Fig. 3e, Supplementary Fig. 3d**) according to the HPA database, suggesting that antibodies targeted broad classes of proteins without obvious bias.

To test antibodies for additional applications, we performed protein complex enrichment and chromatin immunoprecipitation-sequencing (ChIP-seq) with selected antibodies. Protein complex enrichment results are shown in **Fig. 3f**. For example, Pb51585 against PCCA could pull down PCCA-interacting proteins similar to its previously mapped interactome according to STRING^37^ analysis.

ChIP-seq assays in HepG2 for antibodies against SMRC1, SATB1 and NFIC were carried out following previous studies^38^. A total of 93.9 million sequencing reads were generated, and 53.7% were uniquely mapped to the human reference genome. These reads were further processed and yielded 46,380 peaks, representing 29,441, 3,296, and 13,643 binding sites for SMRC1, SATB1 and NFIC, respectively (**Fig. 3g**). Further validation by comparing to commercial ChIP antibody, or with ChIP-qPCR assays, analyses of binding sites and enriched motifs all confirmed that antibodies against those transcription factors could be applied for ChIP-seq (**Supplementary Fig. 3e-h**).

### ADC Antibody/Target Discovery by Differential Array Screening

ADC selectively eliminates cancer cells by linking a toxin, for example Monomethyl auristatin E (MMAE), with an antibody targeting an internalizing tumor-associated antigen characterized by higher expression in tumors than in normal tissues^39^. To identify candidate targets suitable for ADC, we screened PETAL arrays with normal and tumor cell proteomes (**Fig. 4**). More than 3,000 antibodies out of 15,000 lung membrane proteome-positive antibodies showed a fluorescent intensity fold change of >1.5 between tumor and normal samples (**Fig. 4a**). Four antibodies were screened to show both internalization and indirect cytotoxicity from a selection of 500 antibodies with a fold-change ranging from 1.5 to 5 (**Fig. 4b; Supplementary Fig. 4a**). One antibody, Pb44707, with a signal 2.4-fold higher in tumor tissue, was internalized with a half internalization time of 2.5 hours and IC_50_ of ≤100 pM for cell cytotoxicity in PC9 cells. IP and LC-MS/MS identified ProteinX, a putative cancer stem cell marker, as the most likely target protein (data not shown). Target protein was further verified by siRNA blocking experiments in which ProteinX-targeting siRNA caused the loss of surface fluorescent signal in FACS (**Fig. 4d and e**). The cellular binding affinity (EC_50_) of Pb44707 with PC9 was 832 pM (**Supplementary Fig. 4b**) as determined by antibody titration using FACS^40^.

**Figure 4.**
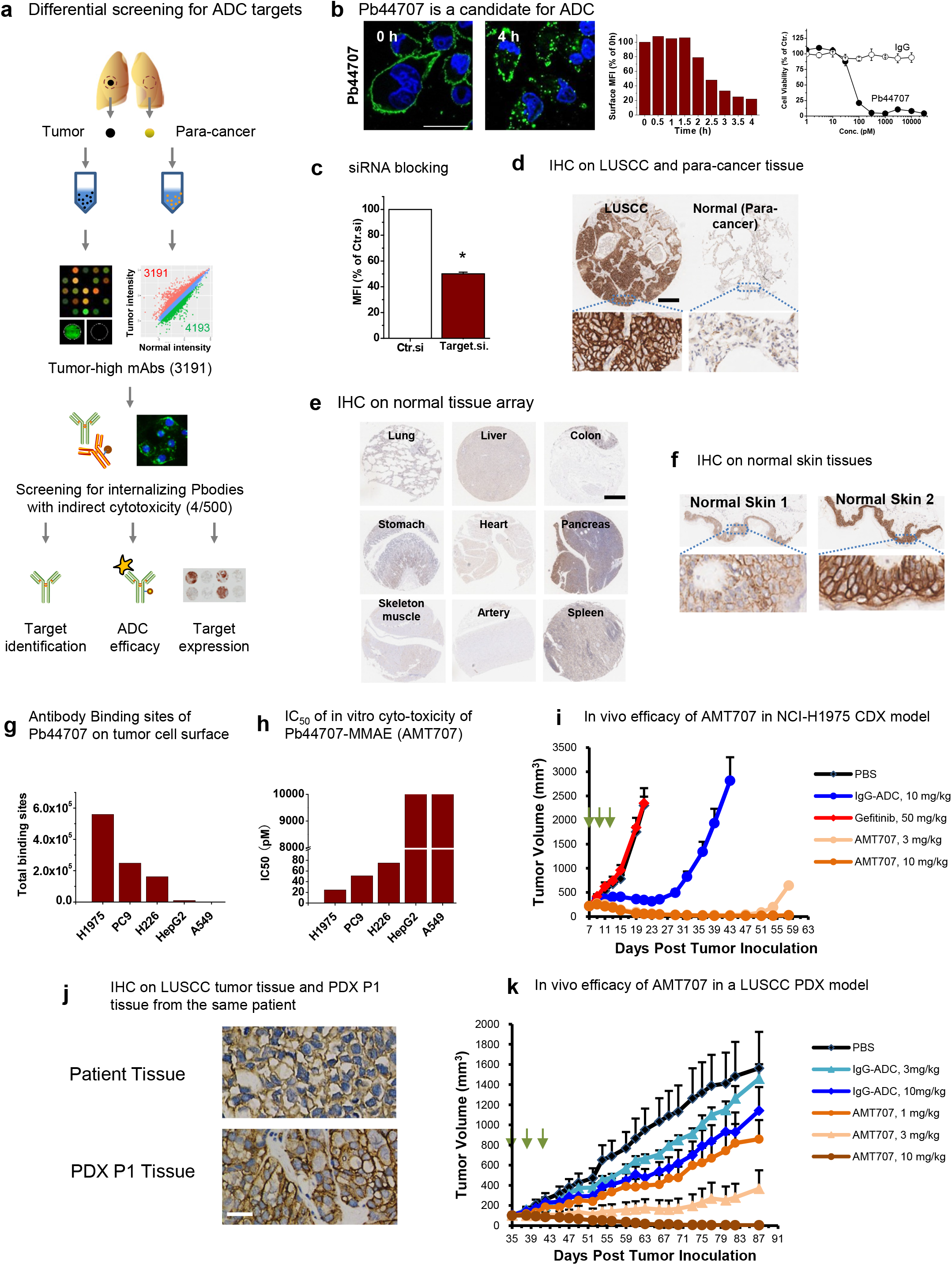
Differential array screening for ADC therapeutic target/antibody discovery. (a) Differential antibody screening for ADC targets. Inset MA scatter plot shows differential distribution of antibody signal intensity of NSCLC and normal lung. N = 1. More than 3,000 tumor-high antibodies were identified. (b) An antibody candidate, Pb44707, for ADC. Antibody ID was labeled on the left of the IF image. IF (0 and 4 hours) image (antibody labeled green and DAPI labeled blue) time course of normalized surface fluorescence in FACS and cell cytotoxicity data is shown. Internalization half time (t_12_) and mean percent growth inhibition ± SEM (n = 3) of the antibody is labeled. IF scale bar = 50 μm. (c) siRNA targeting ProteinX specifically decreases the FACS signal of Pb44707 compared to control siRNA. Normalized mean fluorescent intensity (MFI) ± SEM of PC9 cells as detected by Pb44707. (d) Representative images of the immunohistochemistry staining of Pb44707 in lung squamous cell carcinoma (LUSCC) tumor tissue and para-cancerous tissue from 1 patient. Scale bar = 300 μm. (e) Images of the immunohistochemistry staining of Pb44707 in representative vital normal human tissues. No expression of ProteinX is detected in these tissues. Scale bar = 400 μm. (f) Images of the immunohistochemistry staining of Pb44707 in normal human skin tissues from two healthy persons. Scale bar = 400 μm. (g) Quantification of total binding sites of Pb44707 on the plasma membrane of a variety of tumor cell lines. (h) IC_50_ of Pb44707-ADC on tumor cell lines. N = 1-3 for different cell lines, respectively. (i) Growth curves of the NCI-H1975 CDX tumors of different treatment groups (n = 7/group). Treatment with AMT707, control ADC, or gefitinib (intraperitoneal injection, dosing once a day) was initiated 7 days after tumor inoculation and administered as indicated by arrows. (j) Representative images of the immunohistochemistry staining of Pb44707 in a LUSCC patient tumor tissue (top) and passage 1 (P1) PDX tumor tissue derived from the same patient. Scale bar = 50 μm (k) Growth curves of the PDX tumors of different treatment groups (n = 6/group). Treatment with AMT707 or control ADC was initiated 35 days after tumor inoculation and administered as indicated by arrows.

To evaluate the expression of ProteinX in tumor and normal tissues, Pb44707 was used in immunohistochemistry (IHC) to probe tissue arrays. ProteinX was markedly overexpressed in 60% of NSCLC patients, and more strikingly, in close to 90% of Squamous Cell Carcinoma (LUSCC) (**Fig. 4d, Supplementary Fig. 4c-d**). Furthermore, mRNA of ProteinX showed higher expression in all stages of LUSCC (**Supplementary Fig. 4e**). Most normal tissues were negatively stained with Pb44707 except in skin (**Fig. 4e, and f**), which showed both high and low expression in the population (**Supplementary Fig. 4f**). To build Pb44707 into an ADC molecule, Pb44707 was conjugated to vc-MMAE with an average drug-antibody-ratio (DAR) of 4.23. Pb44707-MMAE (AMT707) demonstrated *in vitro* cytotoxicity in lung cancer cell lines, and its potency was correlated with the ProteinX expression level in these cells (**Fig. 4g and h**). The *in vivo* efficacy of AMT707 was tested in both the cell line-derived xenograft (CDX) model and patient-derived xenograft (PDX) model. For the xenograft model with the gefitinib-resistant NSCLC cell line NCI-H1975 (**Figure 4i**), AMT707 administered at 3 and 10 mg/kg suppressed tumor growth completely until days 47 and 57, respectively, whereas for treatment controls, i.e., PBS, gefitinib and control ADC, there was no or a much lower effect. In a ProteinX-positive LUSCC PDX model (**Fig. 4j and k**), AMT707 haltered tumor growth in a dose-dependent manner, and 10 mg/kg AMT707 suppressed tumor growth completely. Thus, by array screening, we identified multiple endocytic cell surface targets and at least one antibody and its target with a potential to build a lead ADC molecule.

## DISCUSSION

To enable proteome-scale monoclonal antibody development, we constructed PETAL, a hybridoma library consisting of 62,208 mAbs and its corresponding antibody array, the largest antibody microarray reported so far. Taking advantage of antibody multispecificity, PETAL may harbor binders for a large number of proteins in nature^23–26^. Combining PETAL with the global screening capability of antibody microarrays, this platform enables mAb generation at a fraction of the current time and cost. High affinity and desirable specificity of selected mAbs was ensured by a fit-for-purpose workflow. We have demonstrated initial application of PETAL in large-scale antibody generation, affinity proteomics and therapeutic target discovery.

With its capacity to target a large number of proteins in a proteome (**Fig. 2**), our technology provides a solution for proteome/sub-proteome-scale generation of antibodies^41^. We demonstrated that antibodies targeting cell surface and nuclear proteins of cancer and immune cells were efficiently identified. Many of these antibodies were well-suited for IF and IP/ChIP. From an input of 1,818 antibodies, ~200 antibodies capable of immunoblotting/IF/FACS/IP targeting 149 independent membrane and nuclear proteins were identified. Given the input/target ratio of 12:1 (1,818/149), 10,000-20,000 array-positive antibodies would yield more than 1,000 targets.

Antibody-based functional proteomics has not been available for nonhuman species. Here, we demonstrated that broad proteome samples including plants, animals and insects tested so far have all identified binding antibodies corresponding to a large number of protein targets for each proteome. PETAL immediately provided immunoblotting mAbs for initial characterization of proteins in organisms with genomic sequencing information by proteome sample screening and target identification described here. Thus, identified proteome-specific mAb-protein pairs were further used to probe samples from distinctive phenotypes to discover phenotype-specific mAb-target pairs. As demonstrated here, both new and previously reported phenotype-specific targets were discovered for zebrafish heart regeneration and maize seed development. For each protein target, mAbs capable of immunoblotting and IP were obtained. The overall success rate of antibody-target deconvolution was similar to that of human membrane/nuclear screening. Thus, our technology provides a straightforward and productive path to carry out affinity proteomics for numerous proteomes currently without available affinity reagents.

When used for differential profiling of cell membrane proteomes, PETAL provides a “fit-for-purpose” approach for therapeutic target discovery. PETAL relishes the full potential of antibody-based target discovery compared to that of other functional genomics (RNAi or CRIPSR) or MS-based approaches because it delivered the target and lead antibodies at the same time. As demonstrated in this study, differential array screening comparing tumor (NSCLC) with normal lung tissue membrane proteomes identified ProteinX-targeting antibody for building ADC molecule AMT707. More ADC candidate antibodies/targets are expected to be discovered when all 3,000 “tumor high” antibodies are subjected to the screening process.

There are some limitations of the PETAL strategy. The construction of PETAL was a significant investment of time and effort that is difficult to repeat. However, PETAL is now a premade resource that can be readily accessed by researchers to generate mAbs for specific antigens or proteomes or to differentially screen protein samples to identify phenotype-specific protein targets (**Supplementary Fig. 5**). Even though PETAL was made as an antipeptide library, it is likely that other types of antigens could produce equally useful antibody libraries^6^. It is our hope that this work will not only serve as an immediate resource but also stimulate new hybridoma libraries for even more applications. For this purpose, our established industrial-scale monoclonal antibody development capability can be taken advantage of to efficiently build hybridoma libraries/arrays on the scale of tens of thousands of mAbs. Current PETAL only yielded high affinity mAbs for 20-30% of antigens, and this could be improved by increasing the size of the library, which in turn may also increase the success rate of application screening and target identification of PETAL mAbs.

Taken together, PETAL represents a significant improvement over previous antibody array and library approaches. A workflow for array screening and antibody-target deconvolution has been established to eliminate high cost and the long development time of previous methods. We expect PETAL to accelerate functional proteomics by enabling proteome-scale antibody generation and target profiling. We believe PETAL will stimulate the preparation of other antibody libraries; thus, the strategy could be adopted and explored by many other researchers.

## Methods

### Peptide antigen selection

A total of 15,199 peptide antigens, called Proteome Epitope Tags or PETs, were designed from 3,694 proteins representing 418 proteomes. Within each proteome, PETs were selected from unique regions of protein sequence using the heuristic blastp algorithms optimal for short peptide sequence comparison^1, 2^. Peptide antigens representative of predicted surface epitopes from a protein sequence were selected^3, 4^. Peptides were mostly 10-12 amino acids in length to contain 2-3 potential antibody epitopes^5^. Predicted peptide sequences with secondary structures including alpha-helix and beta-sheet were omitted^6^. Special sequences including transmembrane motif, signal peptide, and post-translational modification motif were also not selected. Only disordered or surface-looped regions were selected. Hydrophobic peptides were not selected, and peptide hydrophilicity was calculated by Hopp and Woods method^7^. Finally, peptides with more than one cysteine in the sequence were omitted to avoid synthesis difficulties.

All the peptide antigens were chemically synthesized by GL Biochem (Shanghai) Ltd. The purity and molecular weight of each peptide was evaluated with HPLC and mass spectrum (MS).

### Diversity analysis of PET library

The diversity of the PET peptide library (15,199 peptides, see **Table S1** for detail) is evaluated by comparing the sequence similarity of all peptides against each other. The sequence identity (%) between a peptide and its closest homologs within the library was recorded. PET sequence similarity to two random peptide libraries generated computationally was also compared. The first library was a collection of 15,199 peptide sequences randomly sampled from all species without considering amino acid preference in different species^8^. The second library was constructed by random sampling the entire human proteome (all consensus coding sequences).

### Construction of PETAL monoclonal antibody library

Monoclonal antibodies were developed using a large-scale monoclonal antibody development operation modeled after an assembly line. In the antibody assembly line, each of the close to 100 highly trained technicians performs 1-3 discrete steps (for example, plating fusion cells onto 96-well plates or cell transfer from 96- to 384-well plates) for making hybridomas. An internally built informatics and data system (Antibody Assembler) is used for tracking materials and project status. More than 90% of all materials used are bar-coded to minimize hand labeling. Many steps have automatic data analysis and decision making (for example, clone picking). Together, Antibody Assembly Line is scalable and cost-efficient. PETAL is a premade library built by this highly efficient process. Following traditional hybridoma protocol^9^, after the immunization and fusion, a series of ELISA screens were performed to ensure only the hybridoma clones with the highest affinity to peptide antigens were selected. IgG mAbs were selected using a Sigma antibody isotyping kit (#11493027001). Four to six IgG hybridomas per peptide antigen were selected for multiple rounds of limited dilution subcloning to ensure stability and monoclonality. Each hybridoma cell line was used to prepare milliliters of ascites containing 1-10 mg of mouse IgGs. Mouse strains used for immunization and ascite production were BALB/c and F1 from Shanghai Super-B&K laboratory animal Corp. Ltd. The procedures for care and use of animals were approved by the Abmart Institutional Animal Care and Use Committee.

### Hybridoma V-region sequencing

A mouse IgG primer set from Novagen (#69831-3) was used to amplify the IgG V_H_ and V_L_ regions from selected hybridoma clones. In brief, 1×10^6^ cells were collected for each cell line. Total RNA was extracted using TRIzol reagent (Thermo Fisher, #15596026). The first-strand cDNA was amplified using PrimeScript RT-PCR kit from Takara (#RR104A). PCR products with the expected size (~400 bp for V_H_, and 360 bp for V_L_) were sequenced. The sequences of the PCR products were analyzed by IMGT/V-Quest (http://www.imgt.org)^10, 11^ to define the V_H_ or V_L_ regions and the corresponding subelements. The uniqueness of antibody sequences was evaluated by comparing full-length V (V_H_ and V_L_), frame or CDR sequences using clustal algorithm. The homology matrixes were shown in the heat-map format, and that of the combined CDR sequences was shown in **Supplementary Fig. 1c** as an example.

### PETAL array construction and quality evaluation

Ascites of 62,208 PETAL mAbs were prepared in 162 384-well plates and printed onto nitrocellulose-coated slides (FAST™, Maine Manufacturing, #10486111) in a high-density microarray format (named as PETAL array) using the Marathon system (Arrayjet Ltd, UK). Approximately 100 pL of ascites was printed for each antibody per spot. The array and block layout are shown in **Figure 1c**. A total of 110 blocks were aligned into 10 rows and 11 columns. Each block contains a subarray of 48 x 12 = 576 individual antibody spots, except the subarrays in the last row were printed with 40 x 12 = 480 (3 blocks) or 39 x 12 = 368 (8 blocks) spots. Additional control rows, including a positioning fluorescent spot (Cy3) and a biotin-BSA gradient (0.4-50 pg) of 8 spots, were also printed for each block similar to previous antibody arrays^12, 13^. Biotin-BSA was prepared by saturated labeling of BSA with Thermo Fisher Sulfo-NHS-LC-Biotin (#21336) labeling reagent. PETAL arrays were stored at −80 °C.

To evaluate PETAL array quality, the slides were blotted directly with a mixture of a Streptavidin-Cy3 (Sigma, #S6402) and a Cy5-labeled goat antimouse IgG (Jackson ImmunoResearch, #115-175-146), both at 1:3000 dilution in 1x PBS. Fluorescence of Cy3 and Cy5 were recorded using 532 nm and 635 nm channels by a GenePix 4200A Microarray Scanner (Molecular Devices, LLC.). Images were analyzed by GenePix Pro 6.0 software to give fluorescent intensities of each spot and its corresponding background. Missing or distorted spots, typically controlled under 5% of the total spots, were automatically marked by the software.

Reproducibility of array experiments was evaluated by incubating a triplicate of the same sample with three PETAL arrays. The fluorescent intensity of each spot was normalized using biotin-labeled BSA signal in each block and array. The normalized fluorescent intensities were plotted between every experimental pair. Pearson product-moment correlation coefficient value (R value) was calculated for each data pair in which the r value of +1 means total positive correlation and 0 is no correlation^14^.

### Protein sample preparation

Cell lines, i.e., A431, A549, HEK293T (293T), H1975, H226, Hela, HepG2, HL60, HUVEC, Jurkat, K562, MCF7, PC3, PC9, THP1 and U937, were purchased from ATCC or Stem Cell Bank, Chinese Academy of Sciences (Shanghai, China). Cells were grown or maintained in DMEM or RPMI1640 media following the ATCC cell culture guide. When cells were grown to ~80% confluence, they were dissociated from culture plates by treatment with 1x PBS and 1 mM EDTA for 10-20 minutes. Trypsin was not used to avoid damages on cell surface proteins. Membrane or nuclear fractions of cell lysates were then prepared as described previously^15^. For whole cell lysis, 1x PBS containing 1% NP40, 5 mM EDTA, and protease inhibitor cocktail (Calbiochem, #539134) was added to cells directly and incubated on ice for 30 min. An ultrasonication step was performed before collecting the supernatant. The prepared membrane fraction, nuclear fraction and whole cell lysate were labeled following the cell lines named as MEM, NUC and WCL, respectively. The enrichments of marker proteins in MEM, NUC, and WCL fractions were evaluated with anti-ATP5B (Abmart, #M40013), Histone3.1 (Abmart, #P30266), and beta-Tubulin (Abmart, #M30109) mAbs, respectively.

For tissue samples from plants or animals, whole cell lysates were prepared following a protocol described previously^16^. In brief, frozen tissues were powdered in liquid nitrogen with a pestle, suspended in 10 mL/3 g of tissue extract protein extraction buffer (10 mM Tris, pH 8.0, 100 mM EDTA, 50 mM Borax, 50 mM Vitamin C, 1% Triton X-100, 2%-Mercaptoethanol, 30% sucrose) and incubated for 10 min. An equal volume of Tris-HCl pH 7.5-saturated phenol was then added and vortex mixed for 10 min at room temperature. The phenolic phase separated by centrifugation was recovered and re-extracted twice with 10 mL of extraction buffer. Proteins in the final phenolic phase were precipitated overnight at −20°C with 5x volumes of saturated ammonium acetate in methanol. Protein pellets collected by centrifugation were washed twice with ice-cold methanol and once with ice-cold acetone. Pellets were then dried and dissolved with 500 mM triethylammonium bicarbonate containing 0.5% SDS, pH 8.5. Bacteria lysates were prepared using an ultrasonic apparatus.

### Patient samples

All tissue microarray chips were purchased from Shanghai Outdo Biotech Co., Ltd. Tumor and para-cancerous tissues (normal) were freshly excised from a patient with Non-Small-Cell Lung Carcinoma (**NSCLC**) undergoing surgery. Tumor tissue and matched para-cancerous tissue were homogenized^17^. Briefly, the specimens were cut into 0.5 mm sections before digestion with 0.1% collagenase IV (Gibco, #17104019) for 1 hour at 37°C. The cells were then passed through a 70 μm cell strainer (BD, #352350) and collected by centrifugation for 15 min at 400 g. Plasma membrane proteome extracts were prepared from single cell suspensions of tissues.

### Screening PETAL array with recombinant protein antigens

Recombinant protein antigens were first labeled with biotin using EZ-Link™ NHS-LC-Biotin reagent (Thermo Fisher, #21366) and then hybridized with the PETAL array. Array-bound proteins were incubated with Streptavidin-Cy3. The fluorescent intensity of mAb spots was then recorded by GenePix 4200A Microarray Scanner. Array-positive spots were defined as (signal-background)/background > 3. Protein-binding PETAL mAbs selected from array experiments were further screened through protein-mAb ELISA using a detection limit of 1 μg/mL of protein antigen.

### Screening PETAL array with proteomic antigens

Proteomic antigens including membrane, nuclear, or whole cell lysates were labeled with biotin using EZ-Link™ NHS-LC-Biotin reagent and incubated with the PETAL arrays. Following a similar procedure as described above, antibody spots positive in three independent experiments were then ranked by the averaged fluorescence intensities. A limited number of array positive antibodies (1,000-2,000 according to the expected output of each screening) with high (>10,000), medium (2,000-10,000) and low (500-2,000) fluorescent intensity were selected as candidate antibodies for further validation assays.

### Immunoblotting assays of PETAL mAbs

For recombinant protein immunoblotting, selected mAbs were used to probe 50, 10, 2, and 0.4 ng of recombinant protein antigens.

For immunoblotting of endogenous human protein samples by PETAL mAbs, cell lines were selected according to the protein expression profile from HPA and Uniprot databases. For membrane or nuclear protein targets, corresponding cellular fractions were prepared for immunoblotting. Typically, 20 μg of protein was loaded onto each lane. Support-positive immunoblotting results were evaluated following the criteria described by Antibodypedia (http://www.antibodypedia.com/text/validation_criteria#western_blot) and HPA^2^. Basically, an antibody was qualified as immunoblotting positive when the size of a single or predominant single band on immunoblotting matched or was within 10% of the predicted antigen molecular weight. In some cases, an immunoblotting-positive conclusion was enhanced when the same predicted protein band was detected in two or more different cell lysates. Some antibodies detected multiple bands with different sizes, but the predicted size protein band was also detected.

### IF, FACS validation

A cell line with target protein expression (HPA data) was selected for IF and FACS assays. Known/predicted subcellular localization of the target protein was also obtained from HPA or Uniprot (**Table S4**). For cell surface protein targets, IF and FACS assays were performed under nonpermeable conditions without detergent in the buffers. For intracellular targets, the permeable condition with 0.1% Triton added to the buffers was used throughout. Antibody binding signal was detected using Alexa Fluor 488 and 594 goat anti-mouse IgG secondary antibodies (Jackson ImmunoResearch, #115-545-003 and #115-585-003). In brief, cells attached on cover slips (IF assays) or suspended in 1x PBS (FACS assays) were first fixed in 4% PFA for 10 min. PFA was then removed, and cells were rinsed three times with 1x PBS. Cells were blocked overnight at 4°C in the blocking buffer (1x PBS containing 10% normal goat serum, 0.1% Triton was added for intracellular targets). After removing the blocking buffer, cells were incubated with primary antibody (dilution in the blocking buffer at 1:100-1000) for 3 hours at room temperature. Cells were rinsed 6 times in 1x PBS before being incubated with fluorescence-labeled secondary antibody (diluted in blocking buffer at 1:500 dilution ratio with 1/10000 Hoechst 33258, Sigma #94403) for 1 hour. Finally, cells were rinsed 3 times with 1x PBS. IF images were recorded with the Nikon confocal system A1Si. The 3D reconstruction of the IF results were performed in ImageJ (NCBI free software). IF staining patterns were compared with HPA data to confirm the subcellular localization of the target proteins. The FACS data were collected using the BD Accuri C6 Plus system. A control sample without primary antibody and another sample with isotype control antibody were used.

### IP and Mass-spectrum assays to identify antibody binding target

Immunoprecipitation (IP) assays were performed using CNBr-activated Sepharose 4B (GE, #17-0430-02) by following the user’s manual. In brief, 200 μg of purified PETAL mAbs was cross-linked to 20 μL of hydrolyzed CNBr beads and used to pull-down target protein from 1 mg of cell membrane or nuclear protein samples. Typically, an excessive amount of antibodies was used. A similar procedure was developed following the instructions described previously^18^ to identify the binding protein targets of the tested antibodies.

Essentially, target identification for immunoblotting-successful (yielding single or predominant single band) mAbs used for IP was done by comparing the silver staining result of the IP product with immunoblotting data on samples before and after IP. Expected size band (matched on silver staining and immunoblotting) was selected for MS analysis. For some mAbs, more than one band on SDS-PAGE was selected for MS identification; several identified protein targets could be the binding target of an antibody. For antibodies that failed in the immunoblotting assay, their IP products separated on silver-stained SDS-PAGE were compared to IP products from other mAbs. One predominant specific band or several stoichiometric specific bands were selected for MS analysis

Once one or more bands were selected for MS, 20 μL of IP product was separated in an SDS-PAGE gel and stained with Coomassie blue R-250. The selected target bands were excised and sent to MS facilities (Instrumental Analysis Center of SJTU, or Biological Mass Spectrometry Facility at Robert Wood Johnson medical school and Rutgers, the state university of New Jersey) for target identification using LC-MS/MS on Thermo Q Exactive HF or Thermo Orbitrap-Velos Pro.

Mascot distiller (version 2.6, Matrix Science) or Protein Discovery software (version 2.2) was used to convert raw to mgf or mzML format for downstream analysis. The LC-MS/MS data were searched against Uniprot human (557,992 proteins) for the human cell culture sample, Uniprot zebrafish (61,675) for zebrafish tissue sample or Uniprot maize (137,106) for corn tissue sample. Enzyme specificity was set as C-terminal to Arg and Lys and allowed for two missed cleavages. Furthermore, ±10 ppm and 0.02 Da (Thermo Q Exactive HF) or 1 Da (Thermo Orbitrap-Velos Pro) were used as tolerance for precursor (MS) and product ions (MS/MS), respectively. Carbamidomethylated cysteine was set as complete modifications. N-Terminal protein acetylation and oxidation of methionine were set as potential modifications. Deamidation at asparagine and glutamine and oxidation at methionine and tryptophan were specified as variable modifications.

To ensure MS data quality, we used a threshold of 20 total identified peptide number or 5 nonredundant peptide number to achieve high confidence of the identified protein target. In analyzing the MS result for the antibody, the identified protein list was first prioritized using the total identified peptide number. Proteins that were identified in multiple different antibodies were excluded. For most antibodies in this study, a unique protein with the highest total identified peptide number and matched protein size detected on the silver staining and immunoblotting was selected as the target protein.

The mass spectrometry proteomics data have been deposited to the ProteomeXchange Consortium via the PRIDE^19^ partner repository with the dataset identifier PXD011629 (Reviewer account details: Username, Password: 7ZqfVOM8).

### Abundance distribution and molecular function analysis

The identified membrane, nuclear and other proteins were from the reference database (Nucleoplasm protein database and Nuclear membrane plus Plasma membrane protein database from Human Proteome Atlas). Expression abundance information of human proteins was obtained from the PaxDb database. Function distributions were clustered using the PANTHER classification system^20^ depending on the molecular function.

### ChIP-Seq assay

The ChIP and input DNA libraries were prepared as previously described^21, 22^. Briefly, 10 million HepG2 cells were cross-linked with 1% formaldehyde for 10 min at room temperature and then quenched with 125 mM glycine. The chromatin was fragmented and then immunoprecipitated with Protein A + G magnetic beads coupled with antibodies against SMRC1, SATB1 and NFIC. After reverse crosslinking, ChIP and input DNA fragments were used for library construction with NEBNext Ultra Ligation Module (E7445, NEB). The DNA libraries were amplified and subjected to deep sequencing with an Illumina sequencer. The ChIP-seq data processing was performed as we reported recently^22^. C/s-regulatory sequence elements that mediate the binding of SMRC1, SATB1 or NFIC were predicted with MEME-ChIP^23^.

### Internalization Assay

For the IF assay, live PC9 cells were cultured on coverslips and incubated with 10 μg/mL of mAbs for 1 hour on ice before being washed 3 times with PBS. Cells were then cultured at 37°C for 0, 2, or 4 hours before fixation with 4% PFA. Antibodies were then labeled with Alexa Fluor 488 conjugated anti-Mouse antibody. Images were acquired by Nikon confocal system A1 Si.

For the FACS assay, live PC9 cells were incubated with 10 μg/mL of mAbs for 0.5 hour on ice before being washed 3 times with PBS. Cells were then cultured at 37°C for up to 4 hour before fixation with 4% PFA. Cells were then stained with Alexa Fluor 488-conjugated anti-Mouse antibody and analyzed with FACS. Surface MFI was calculated. Surface MFI, which represented surface localization of mAbs, was measured by FACS.

### Indirect cytotoxicity assay

PC9 cells were cultured in 96-well plates at 2000/well confluence overnight. Cells were treated with serial dilution of mAbs together with 2 μg/mL of Monomethyl auristatin E (MMAE)-conjugated secondary goat anti-mouse IgG antibody for 72 hours. Cell number was then calculated by Cell Counting Kit-8 (CCK8, Dojindo, #CK04-20). Antibody-drug conjugation services were provided by Levena Biopharma, Nanjing.

### *In vivo* tumor models

For the cell line-derived xenograft (CDX) model, 5×10^6^ NCI-H1975 cells were suspended in Matrigel (BD Biosciences, #354234) and injected subcutaneously (s.c.) to the right flank of female BALB/c nude mice (jsj-lab). For studies with the patient-derived xenograft (PDX) model, the tumor fragments from patients with lung squamous cell carcinoma were passaged twice in NOD-SCID mice (Beijing Vital River Laboratory Animal Technology Co., Ltd.). Tumor fragments obtained from *in vivo* passage were then implanted s.c. in the right flank of female BALB/c nude mice (jsj-lab). Body weight and tumor volume (0.5 x length x width^2^) were measured every 3 days. Mice were randomized into control and treatment groups based on the primary tumor sizes (median tumor volume of approximately 100 mm^3^). Pb44707-ADCs and control ADCs were administered intravenously (i.v.) every third day and repeated for a total of three times (Q3Dx3). Gefitinib (Selleck, ZD1839) was administered intraperitoneally (i.p.) every day.

### siRNA knockdown and overexpression

PC9 was transfected with siRNA targeting human PROTEINX (sense, 5’-CUA CUU UAC UGG AAG GUU Att-3’; and antisense, 5’-UAA CCU UCC AGU AAA GUA Gtt-3’), which has been reported previously^24^ or control siRNA (sense, 5’-UUC UCC GAA CGU GUC ACG Utt-3’; and antisense, 5’-ACG UGA CAC GUU CGG AGA Att-3’) by Lipofectamine 2000 (Thermo Fisher, #11668019) 48 hr before performing experiment.

PIEZO1-GFP plasmid used for overexpression validation was a gift from Prof. David Beech, which was described previously^25^. PIEZO1-GFP was transfected into HUVEC cells by lippofectamine200. Cells were fixed and stained with anti-PIEZO1 antibody following IF procedure described above.

### Peptide blocking assay

1 mg/ml Pb44707 was pre-incubated with 1 mg/ml PROTEINXs recombinant protein (Abcam) or PROTEINX peptide in 1:1 ratio at 4°C overnight before used in FACs analysis.

### Antibody cellular binding site quantification

The antibody binding sites on cell lines were determined with the QIFIKIT (Dako, # K0078) according to the manufacturer’s instructions.

## Supplementary Figure legends

**Supplementary Figure 1.**
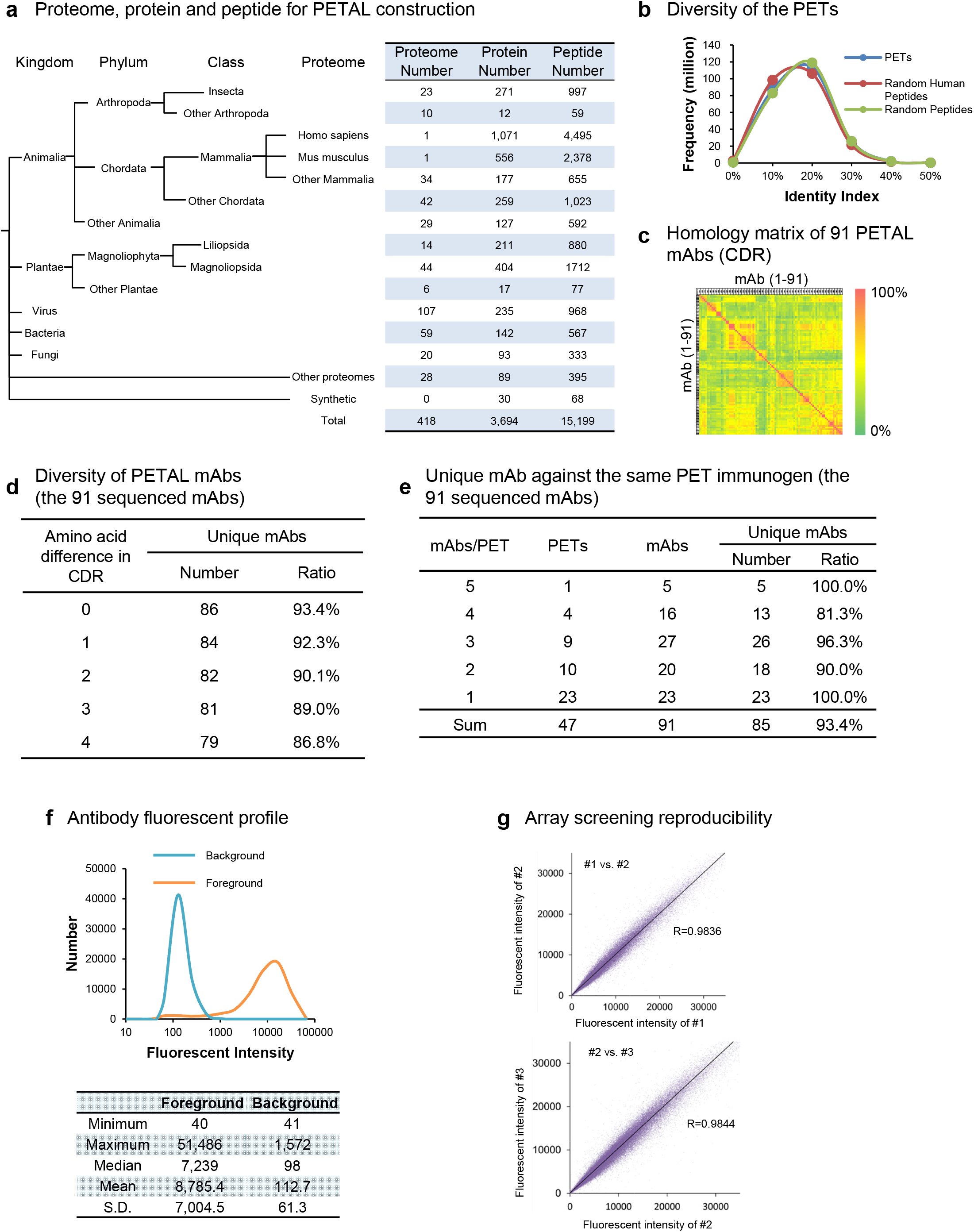
PETAL antigens, antibody diversity and array performance. (a) Proteome, protein and peptide for PETAL construction. Peptide length is 1012 amino acids. Artificially designed peptides, such as tag peptides and random-generated peptides, are labeled Synthetic. The table on the right showed the number of proteomes, proteins and peptides. (b) Diversity of the PET library. PET sequence identity comparison with two randomly created peptide libraries. One from human protein sequences was labeled Random Human Peptides, whereas another from all NCBI sequences of the same length was labeled Random Peptides. All three peptide sets showed a similar sequence identity profile by pairwise alignment, suggesting that the PET library is random and diverse. (c) Homology matrix of the CDR sequences of 91 PETAL mAbs shown as heat map. Homology scale is shown on the right side of the panel, in which red is 100% homology and green is 0% homology. Overall, more than 90% of the sequenced mAb sequences were unique. (d) Diversity of PETAL mAbs. The relationship between the number/percentage of unique PETAL mAb sequences and the number of CDR amino acid differences is shown. (e) Unique mAbs against the same PET immunogens. Overall, more than 90% of mAb sequences against the same peptide antigen were unique, implying that these antibodies likely either recognize different amino acid epitopes or bind to the same epitope with different affinities. (f) Fluorescence profile of printed antibodies with the distribution of foreground and background fluorescent intensities of printed antibodies. (g) Three independent array-hybridization experiments (marked as #1, #2 and #3) were performed using a proteome sample (total protein extract from a human cancer cell line). The signal intensities were then plotted, and the reproducibility, i.e., the coefficient values (R) were calculated among these 3 experiments (labeled as #1 vs. #2 and #2 vs. #3).

**Supplementary Figure 2.**
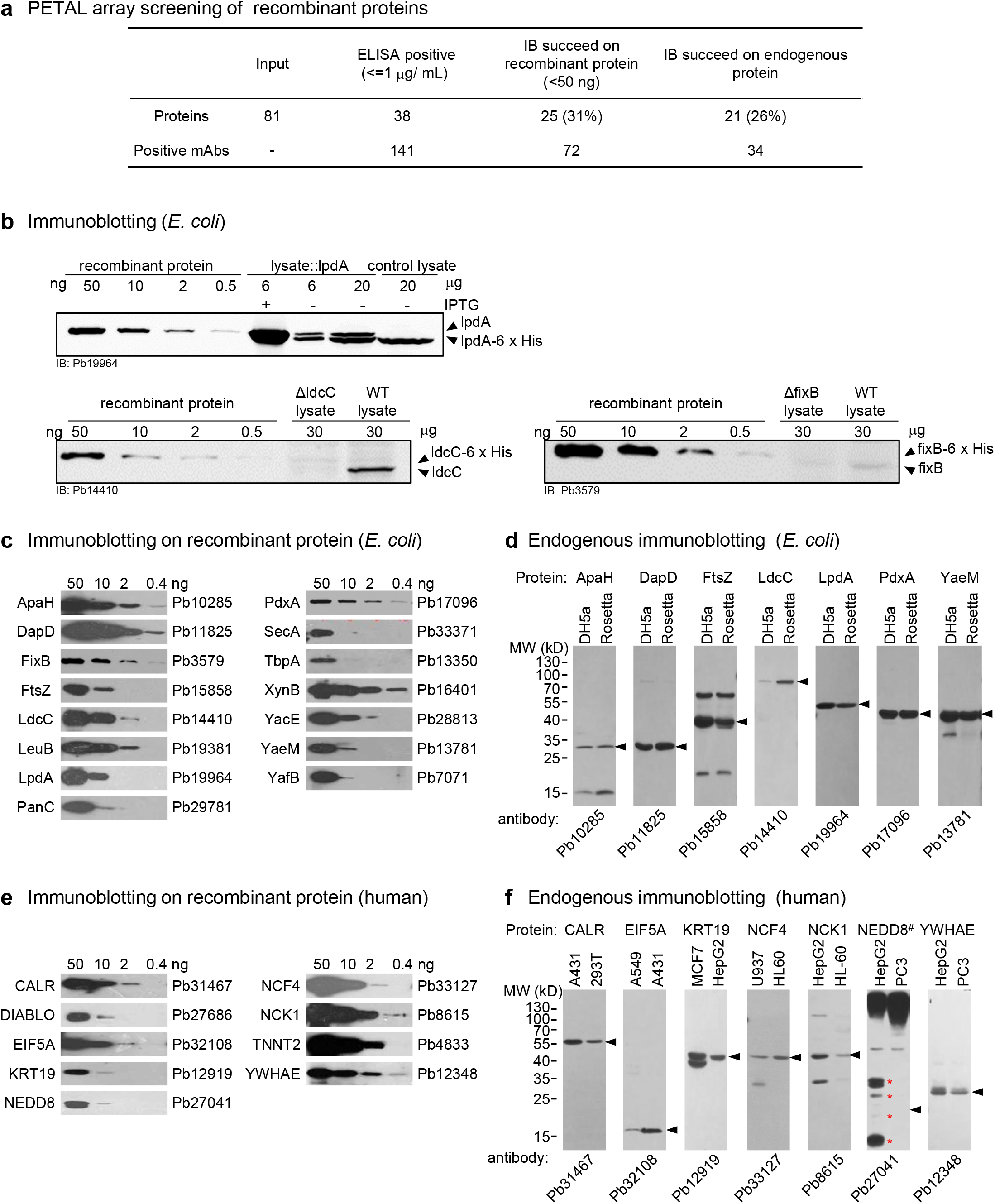
PETAL screen by protein antigens. (a) mAb discovery for recombinant proteins by PETAL array. Approximately 30% of recombinant proteins yielded successful immunoblotting mAbs using recombinant (31%) or endogenous protein samples (26%). (b) (top panel) Immunoblotting (IB) validation of Pb19964 against *E. coli* protein lpdA. Purified recombinant proteins of serial dilutions and total lysate of *E. coli* with lpdA overexpression were immunoblotted with the corresponding antibodies. The total lysate of an *E. coli* strain with the expression vector only was included as control. The arrowheads on the right marked the His-tagged protein and endogenous protein. (middle and bottom panels) Immunoblotting validation of Pb14410 and Pb3579 against *E. coli* proteins ldcC and fixB, respectively. Purified recombinant proteins of serial dilutions and total lysate of *E. coli* from ldcC (ΔldcC), fixB (ΔfixB) knockout strains were immunoblotting tested with the corresponding antibodies. The total lysate of WT *E. coli* strain was included as control. The arrowheads on the right mark the His-tagged protein and endogenous protein. (c) Immunoblotting of antiprotein PETAL mAbs on additional *E. coli* proteins. Protein name is labeled on the left side of each panel. The corresponding PETAL mAbs IDs are labeled on the right. (d) Immunoblotting of antiprotein PETAL mAbs on endogenous cell lysates from *E. coli;* 20 μg of total lysate was loaded on each lane. (e) Immunoblotting of antiprotein PETAL mAbs on additional human recombinant proteins (f) Immunoblotting of antiprotein PETAL mAbs on endogenous cell lysates from human cell lines; 20 μg of total lysate was loaded on each lane. #: NEDD8 is a ubiquitin-like protein; therefore, the ladder-like pattern and aggregation in large molecular weight were judged to be a positive result. Red asterisks marked the predicted position of NEDD8 monomer, dimer, trimer, and tetramer.

**Supplementary Figure 2.**
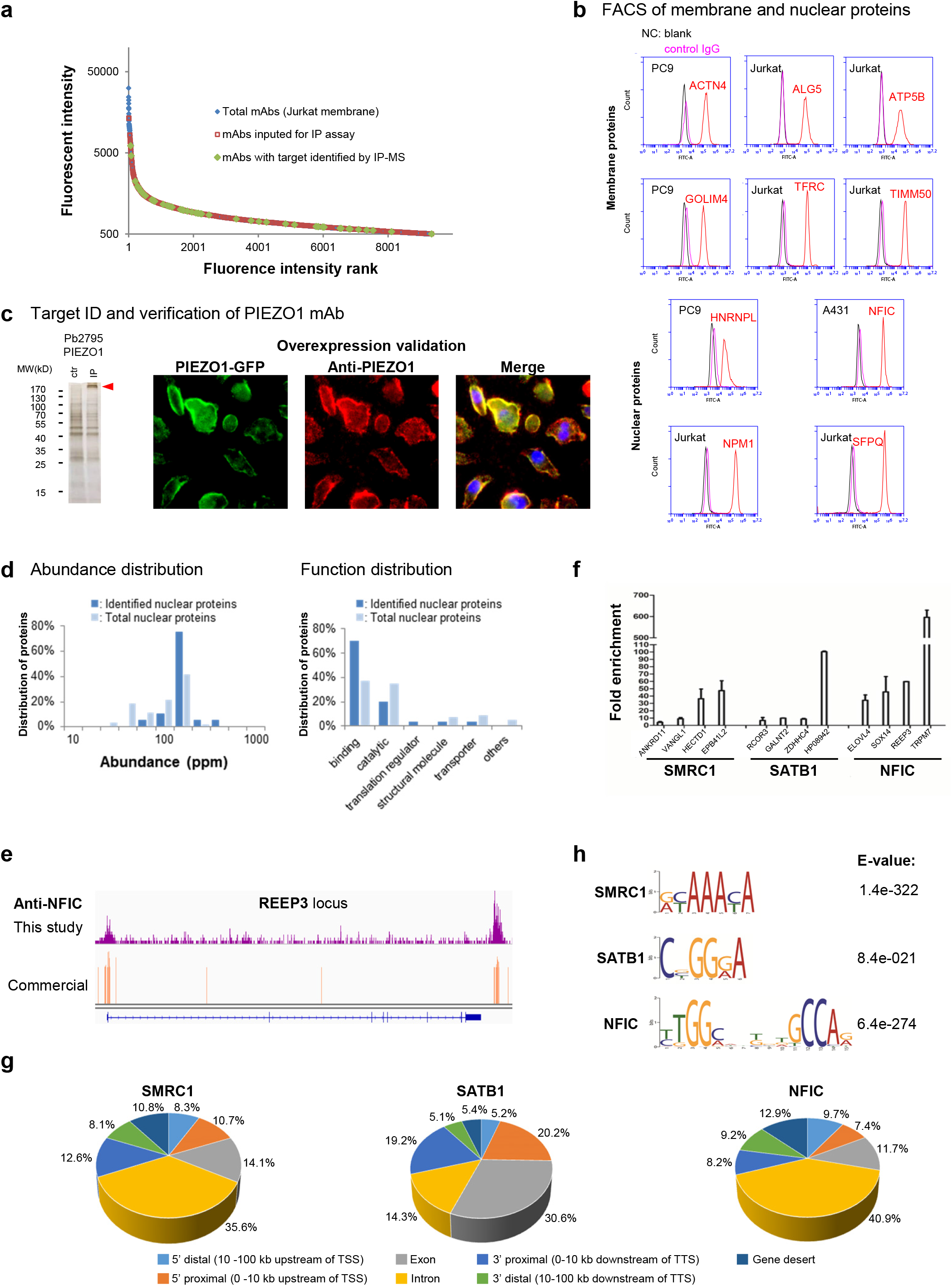
Proteome-scale antibody generation for human membrane and nuclear proteins. (a) Distribution of fluorescent intensity of array-positive antibodies from Jurkat cell membrane proteome screening (blue). Approximately 15,000 antibodies were positive for binding with a fluorescent signal intensity in the range of 500-15,000. Red dots represent antibodies used in the IP assay, and green dots represent those successful for application screening (Immunoblotting/IF/FACS/IP) and target identification, showing similar success rates across the antibody fluorescence intensity scale. (b) FACS analyses for endogenously expressed membrane and nuclear proteins. Figure labels are the same as in **Fig. 3b** for TFRC. (c) Overexpression validation of anti-PIEZO1 antibody Pb2795. Silver staining of Pb2795 IP product (lane IP) is shown on the left side. Lane ctr was loaded with IP product from an irrelevant antibody. Arrowhead marks the dominant IP band of PIEZO1. On the right side, HUVECs were transfected with PIEZO1-GFP (green) expression vector and stained with Pb2795 (red). The result shows fluorescent overlap of GFP and antibody (yellow in the right panel). Cells were stained with DAPI (blue). (d) Similar to **Fig. 4e**, data for nuclear protein targets. The abundance and function distribution of the proteins identified from Jurkat cell nuclear proteome. (e) Snapshot of IGV browser showing ChIP-seq read density at the REEP3 locus. The purple peaks were derived from PETAL NFIC antibody, and the orange ones were generated with a commercial NFIC antibody^42^. Antibody for NFIC identified known palindromic sequence TTGGCNNNNNGCCAA, whereas the commercial antibody did not enrich this sequence. (f) ChIP-qPCR validation performed on four randomly chosen peaks from each ChIP-seq assay of SMRC1, SATB1 or NFIC antibody. The results were normalized to input chromatin, showing all chosen regions were fairly enriched. (g) Genomic location distribution of binding sites of SATB1, SMARC1 and NFIC relative to the nearest transcription units. The percentages of binding sites at the respective locations were shown. (h) Predominant binding motifs of SATB1, SMRC1 and NFIC. These motifs were identified from the ChIP-seq datasets of SATB1, SMRC1 and NFIC, respectively. Known NFIC motif (TTGGCNNNNNGCCAA) was recovered using an antibody for NFIC.

**Supplementary Figure 4.**
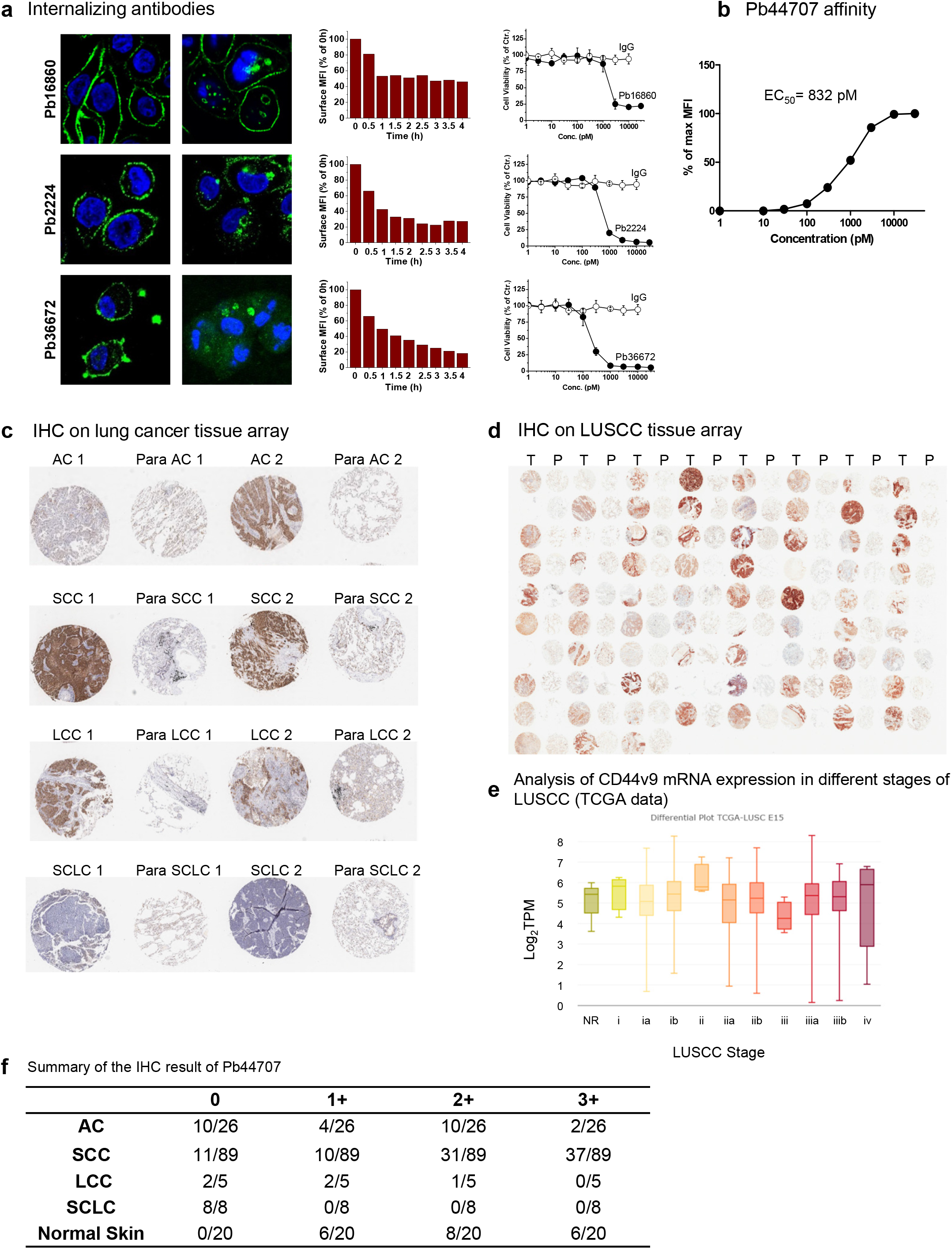
Differential array screening for ADC therapeutic target/antibody. (a) Antibody candidates for ADC. Antibody ID was labeled on the left of the IF image. For each antibody, IF (0 and 4 hours) image (antibodies labeled green and DAPI labeled blue) time course of normalized surface fluorescence in FACS and cell cytotoxicity data is shown. Internalization half time (t_1/2_) and mean percent growth inhibition ± SEM (n = 3) for each antibody is labeled. IF scale bar = 50 μm. (b) Pb44707 cellular binding affinity. Representative image showing the binding affinity of Pb44707 with PC9 cells. The EC_50_ is 832 pM. N = 3. (c) Examples of IHC of Pb44707 on lung cancer tissue array. Subtypes of NSCLC were labeled squamous cell carcinoma (SCC), adenocarcinoma (AC), and large cell carcinoma (LCC). Small cell lung carcinoma was labeled SCLC. For each tumor type, IHC data from two patients and their papa tumor tissues are shown. ProteinX was found overexpressed in 40% of AC and LCC, in 90% of LUSCC and not expressed in SCLC. (d) ProteinX expression in LUSCC. Pb44707 was used to probe a tissue array of LUSCC patients with tumor (labeled as T) and para-cancerous (labeled as P) samples; 85% of LUSCC patients had overexpression of ProteinX. Intensity of ProteinX overexpression on IHC was graded as high (+++, 36%), medium (++, 37%) and low (+, 12%). See (g). (e) ProteinX mRNA expression analyzed at different stages of LUSCC using the TCGA database. (f) Summary of ProteinX expression in NSCLC and skin.

**Supplementary Figure 5.**
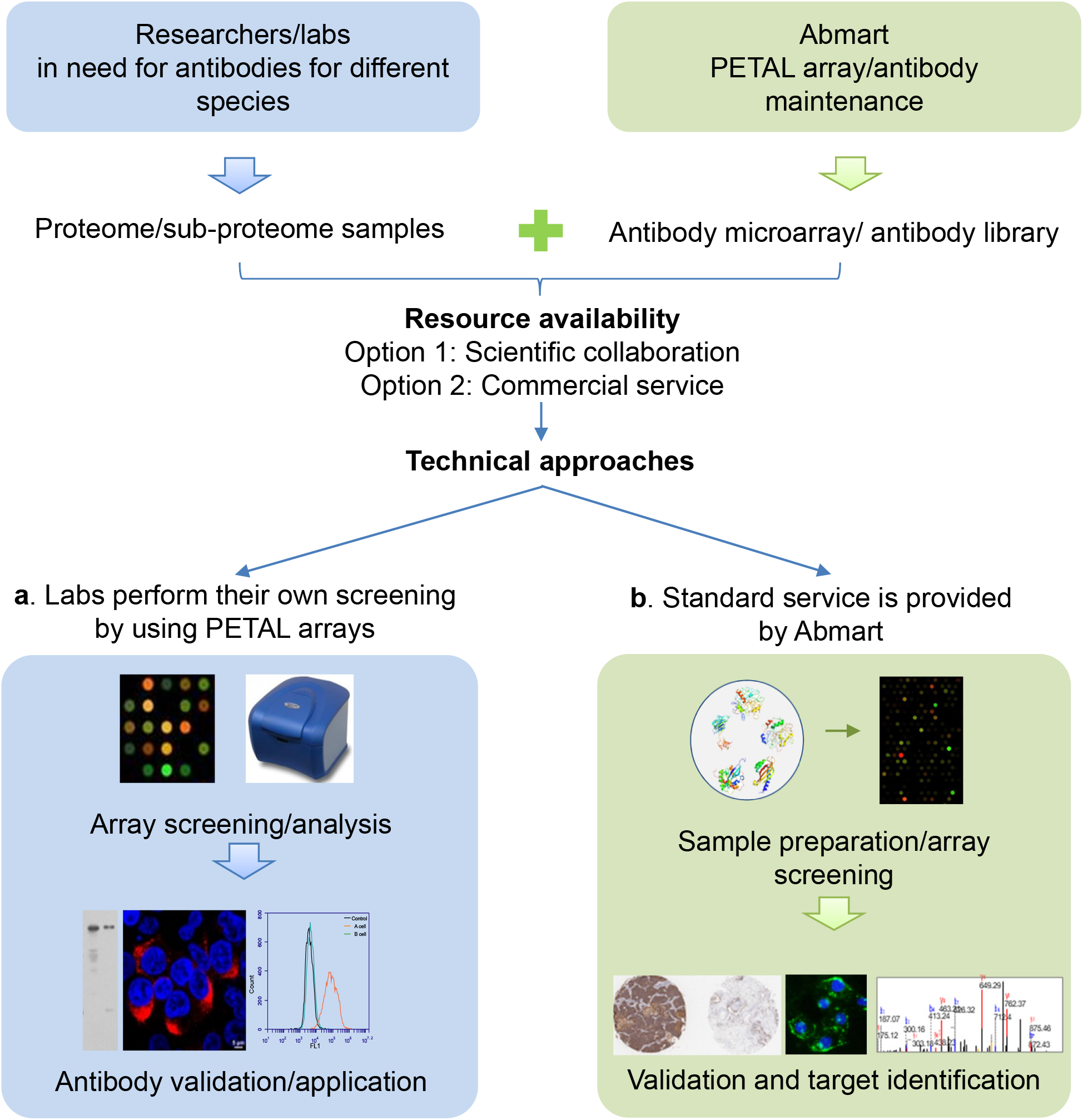
Two approaches to access PETAL library/array. (a) PETAL arrays are provided to labs to run their own screenings. Any lab with access to a microarray facility, especially if equipped with a microarray scanner, is able to perform the microarray experiment by themselves using the PETAL microarrays following the procedure that we described. Once the array screening is performed with antigens of interest, a list of the identified antibodies, or simply the array hybridization result, could be sent back. Antibodies can then be provided at any point dependent upon the researcher’s antibody need. (b) Standard screening service is provided. Tissue samples/total protein lysates or other types of proteome/subproteome samples are provided for screenings following the process described in this report. As the final outcome, scientists will then be provided with a set of validated monoclonal antibodies. In both approaches, arrays/services will be provided by Abmart and third party suppliers through two options: Option 1, scientific collaboration; Option 2, commercial service.

